# Does mother know best? Range-wide narrowing of host preference in *Aricia agestis* confers fitness benefits but may incur long-term costs

**DOI:** 10.1101/2025.09.04.674201

**Authors:** Brooke Zanco, Maaike de Jong, Ebba Widman, Florencia M Camus, Jon Bridle

## Abstract

Biological communities are responding to changes in climate through a combination of range shifts and evolution *in situ*. However, even in systems where we have genomic and phenotypic evidence of recent adaptation, the ecological and fitness consequences of such evolutionary change remain underexplored, especially in terms of consequences for interactions between species. European Lepidoptera are excellent systems for understanding such shifts, given our exceptional knowledge of their ecology, natural history, and past distributions. The UK Brown Argus butterfly (*Aricia agestis*) has expanded rapidly northward in response to climate change. This expansion has been associated with evolutionary shifts in maternal host preference, from the chalkland restricted but locally common perennial Rockrose (*Helianthemum nummularium*) to more broadly distributed, but locally rare annual Geraniums (*Geraniaceae*). Such a rapid range shift offers an excellent opportunity to test how the evolution of novel biotic interactions associated with climate adaptation affects individual fitness and population resilience.

Using common garden assays of host preference on females sampled across the UK in 2013 and 2023, we show for the first time that female preference for *Geranium molle* or Rockrose hosts vary among individuals within as well as among sites and to different extents at different locations. This variation is consistent with genomic signals of recent selective sweeps on these traits. In particular, we show that in 2013, females in the new part of the range are more likely to oviposit on the locally dominant host, while females from the ancestral range will use both hosts, regardless of the locally dominant host. We also demonstrate a significant temporal shift in behaviour, with a narrowing of host preference towards *Geranium molle* recorded in *A.agestis*’ ancestral range in 2023.

To evaluate the fitness consequences of these shifts in biotic interactions, we assessed larval performance of 1412 larvae from 49 families on both host plants. Irrespective of maternal preference, feeding on *Geranium molle* hosts conferred higher survival, faster development and larger adult body size under laboratory conditions. These fitness outcomes are likely linked to *Geranium molle*’s significantly higher protein: carbohydrate ratio when compared to Rockrose. Moreover, we also found that larvae fed *Geranium molle* produced adults with elevated mass-independent resting metabolic rates, a rare demonstration that larval diet has lasting metabolic consequences in a wild butterfly.

Crucially, we found no evidence of trade-offs between preference and performance: even offspring of females that laid exclusively on Rockrose also performed better on *Geranium molle*. These results suggest that *Geranium molle* confers a general performance advantage, enhancing traits likely to promote population persistence where *Geranium molle* is reliably available. However, despite selection favouring *Geranium molle* in warming climates, we also show that a narrowing of host preference towards these more widespread host plants throughout the UK range is likely to make *A.agestis* populations more vulnerable under future warmer, drier conditions, when compared to Rockrose. Together these findings highlight a central challenge in evolutionary biology: what happens when traits that are adaptive in the short term spread rapidly through populations, but are likely to become maladaptive as environmental conditions continue to change?

## Introduction

Climate change is reshaping ecological systems at a rapid and accelerating pace, altering species’ distributions, changing biotic interactions, and imposing novel selection pressures (Hoffmann and Sgró, 2011; Pecl *et al*., 2017; Bridle and Hoffmann, 2022; Lawlor *et al*., 2024). These changes are widespread but highly variable across taxa and ecosystems, making predictions of species’ persistence or decline challenging (Scheffers *et al*., 2016). Some groups of organisms are particularly sensitive to these shifts. For example, herbivorous insects, whose life cycles are tightly linked to the phenology and chemistry of their host plants, are highly vulnerable to changes in plant availability and quality (Merrill *et al*., 2009; Pateman *et al*., 2012). As temperatures rise and precipitation patterns shift, the abundance and quality of the resources on which these insects depend are changing in ways that require rapid ecological and evolutionary responses (Stewart, 2020; Stewart *et al*., 2021; Brodie *et al*., 2025). In these systems, shifts in host plant use represent one of the most ecologically significant forms of adaptation, enabling herbivorous insects to persist in novel environments or track shifting climatic envelopes (Bridle *et al*., 2013; Buckley and Bridle, 2014; Stewart *et al*., 2021). Such shifts are particularly important given recent projections suggest that range shifts for many plant species will lag behind present-day rates of climate change, thus, if species are unable to shift host use, they could risk extinction (Iverson, Schwartz and Prasad, 2004; Scheller and Mladenoff, 2008; Sharma *et al*., 2022).

Host shifts can arise through a range of mechanisms, including phenotypic plasticity, genetic change, or an interaction between both (Vertacnik and Linnen, 2017; Saastamoinen *et al*., 2018; Verspagen *et al*., 2020; Singer and Parmesan, 2021; Stewart *et al*., 2021). Although these shifts may allow range expansion and short-term persistence, they can be associated with increased specialisation, ecological mismatch, or dependence on unstable resources (Schlaepfer, Runge and Sherman, 2002; Bridle *et al*., 2013; Buckley and Bridle, 2014; Hale, Treml and Swearer, 2015). Despite this, the long-term consequences of host shifts in environments that are rapidly changing remain poorly understood, particularly in specialist species where such obligate biotic interactions are constrained by evolved physiological and behavioural interactions (Ehrlich and Raven, 1964; Janz and Nylin, 1998; Slove and Janz, 2011). Consequently, quantifying variation in host preference within and among populations is crucial for understanding evolutionary potential and the potential consequences of host shifts under environmental change.

In specialist herbivores, host shifts often require substantial adaptive changes, especially when the new host is chemically distinct from the original host. Such phenotypic changes include modifications to detoxification pathways, sensory systems, as well as digestive enzymes and forms of parental provisioning and microclimate use (Després, David and Gallet, 2007; Bass *et al*., 2013; Vertacnik and Linnen, 2017; Dar *et al*., 2024). These adaptations typically carry trade-offs: improved performance on a novel host may reduce the ability to utilise ancestral hosts. Such trade-offs can occur through antagonistic pleiotropy, where alleles that improve performance on the new host directly reduce performance on the old host. Trade-offs may also occur through mutation accumulation, where alleles that were neutral or beneficial on the ancestral host accumulate deleterious mutations because they are no longer under selection (Grosman *et al*., 2015). Moreover, maintaining preference for multiple hosts may itself be costly, requiring broader sensory and regulatory capacity that might reduce oviposition precision or efficiency (Futuyma and Moreno, 1988; Bernays and Wcislo, 1994).

The Brown Argus butterfly (*Aricia agestis*) is a compelling system in which to examine these dynamics. In the UK, *A. agestis* has expanded its range northward over recent decades (Asher *et al*., 2001), as climate change makes new habitats thermally suitable (Warren *et al*., 2001; Pateman *et al*., 2012). In long-established parts of the range, *A. agestis* is mostly restricted to chalk/limestone grassland dominated by the perennial host plant, Rockrose (Cistaceae, *Helianthemum nummularium*). However, some populations persist at locally warm sites dominated by host plants in the Geraniaceae family (*Geranium molle, Geranium dissectum* and *Erodium cicutarium*), short lived annuals, which are not closely related to Rockrose (Soltis *et al*., 2011), and highly variable to climate-driven phenological variation (Bridle *et al*., 2013; Buckley and Bridle, 2014; Stewart, 2020; Stewart *et al*., 2021). Extensive transplant experiments have shown that populations in the novel part of the range have narrowed their host preference towards these Geraniaceae species (Buckley and Bridle, 2014). This narrowing of preference is likely genetically mediated, with genomic analyses revealing strong selection on host-use loci across the species’ range (Bridle *et al*., 2013; de Jong *et al*., 2023). Although much research has focused on the effects of narrowing host preference in populations at the expansion front, evidence suggests that host preference in *A. agestis* is also evolving quickly within long-established populations (de Jong *et al*., 2023), where persistence at a given location under continued climate warming presumably necessitates rapid *in situ* evolution (Bridle and Hoffmann, 2022). This phenomenon raises important questions about whether traits selected at the range edge might be expected to have emerged from, or spread into, long established populations, and about what ecological or evolutionary costs may emerge (Thomas *et al*., 2001; Lenoir and Svenning, 2015; Saastamoinen *et al*., 2018)..

Although rapid adaptation increases short-term fitness, it may also carry long-term risks. In particular, strong directional selection can reduce genetic variation and push populations toward narrower ecological niches (Hoffmann and Bridle, 2022; Chevin and Bridle, 2025). In herbivores like *A. agestis*, where female oviposition decisions are closely linked to offspring performance, a shift towards novel hosts that allow growth into new climate zones could undermine long-term persistence if those hosts are nutritionally unstable, or vulnerable to continued climate warming. In the case of *A. agestis*, we already know that *Geranium molle* hosts in the novel parts of the range vary more in condition, availability, and microclimate than Rockrose, while also showing significant variability in phenology, depending on a narrow window of climatic conditions, in particular the availability of rain during the late summer (Stewart, 2020; Stewart *et al*., 2021). What remains unclear, however, is the extent to which this environmental variability translates into differences in plant traits that directly influence larval performance, such as protein availability, photosynthetic capacity and secondary metabolite production.

An often-overlooked consequence of host preference shifts is how larval diet shapes resting mass-independent metabolic rate. Diet not only affects growth directly, but also influences the costs of digestion and nutrient absorption (Roces and Lighton, 1995; Nespolo, Castañeda and Roff, 2005; Naya *et al*., 2007) and the development of energy-demanding tissues such as the fat body and flight muscles, which together determine adult metabolic rate (Yang and Joern, 1994; Desai and Hales, 1997; Roark, Bjorndal and Bolten, 2009). For example, in honeybees, the protein-to-carbohydrate balance of larval diets has been shown to alter the scaling of resting metabolic rate, linking early nutrition to differences in adult energy use (Nicholls, Rossi and Niven, 2021). Despite this finding, no study to date has tested the effects of host preference shifts in a range expanding species on resting mass independent metabolic rate.

The preference–performance hypothesis predicts that selection should favour mothers that lay eggs on hosts that maximise offspring fitness (Thompson, 1988). However, empirical studies frequently reveal mismatches between preference and performance, particularly during or after host shifts (Gripenberg *et al*., 2010; Singer, 2015). Host preference can in such cases be driven by ecological factors such as inter- and intraspecific competition with other larvae (Friberg and Wiklund, 2009), or adult diet and preferred microclimatic conditions (Clark, Hartley and Johnson, 2011; Henry, Bocedi and Travis, 2013). However, whether preference–performance alignment is constraining or facilitating adaptation in *A. agestis* remains unclear. A key possibility is that maternal oviposition choices align with offspring performance, in which case shifts in preference would directly facilitate host-use change (García-Robledo and Horvitz, 2012), whereas a mismatch between choice and offspring success would point to hidden trade-offs (for example, benefits to mothers from choosing a particular microhabitat or host that reduce offspring fitness) (Clark, Hartley and Johnson, 2011; Murphy and Loewy, 2015). Distinguishing these scenarios is central to understanding whether temporal shifts in preference represent adaptive tracking of the best host, or are constrained by maternal trade-offs that produce lagged or complex evolutionary responses (Warren *et al*., 2001; Singer and Parmesan, 2018; Hoffmann and Bridle, 2022).

In this study, we present the first data on among individual and among site variation in maternal host preference across the UK in 2013 and 2023 (20 generations apart, as *A.agestis* is bivoltine), in common garden assays where individual females have been given a choice of *Geranium molle* or Rockrose experimental host plants. We then go on to assay reciprocal larval performance, including measurements of developmental rate, body size, resting metabolic rate, in relation to maternal preference, within and among families, combined with host plant trait analyses. Using these methods we test for: (a) variation within and among populations in the established versus novel parts of the range, and between 2013 and 2023; and (b) variation in larval performance, adult fitness and metabolic rate when reared on these different host plant species under common garden conditions, including where maternal choice of host plant is circumvented. (c) macronutrient availability of host plants grown under common conditions without stress; and (d) how host plant quality is altered under experimental temperature and drought stress. This integrated approach allowed us to test for shifts in host plant use, and therefore adaptive potential, in *A. agestis*. It also enabled us to evaluate how maternal preference aligns with offspring performance, and whether potential trade-offs emerge when host plant use shifts. Finally, we consider whether increasing specialisation on *Geranium molle* might constrain future population persistence. Together, these analyses provide some of the first empirical evidence for how insect–host plant interactions evolve under climate change, and for the extent to which short-term adaptation may come at the cost of long-term resilience.

## Materials and Methods

### Host Preference

In 2013 and 2023, host preference in *A. agestis* was assessed during the second brood (mid-August to early September) using oviposition assays in mesh cages containing greenhouse-grown *Helianthemum nummularium* and *Geranium molle*.

### 2013 Host preference assays across the UK

In 2013 eight *A. agestis* populations (19– 45 females per site) were sampled across most of their latitudinal distribution in the UK during the 2^nd^ generation (late July to late August). Each site was sampled at least twice during the collecting season to test for any effects of early vs late emergence. Sites were chosen to include long established (present since 1970 – 1982) and newly colonised sites (since 1995 – 1997); (Thomas *et al*., 2001; Buckley and Bridle, 2014) (Figure 1c), and sites were classified as either dominated by Geraniaceae (which includes *Geranium molle, Geranium dissectum and Erodium cicutarium*) or Cistaceae (*Helianthemum nummularium (Rockrose)*) (de Jong *et al*., 2023). Individuals were placed into 0.5 × 0.5 m wire mesh cages at random positions in a grid at Fenswood Farm, Long Ashton (Bristol University Experimental Gardens, Bristol, BS9 1JB, UK). Each cage contained four greenhouse-grown plants: two *Helianthemum nummularium* grown from plugs: purchased from Naturescape (Maple Farm, Coach Gap Lane, Langar, Nottinghamshire, NG13 9HP, United Kingdom) and two *G.molle* grown from seed: purchased from Emorsgate Seeds (Manor Farm, Langridge, Bath, BA1 9BX, United Kingdom). All plants were reared under natural light in Bristol Fenswood Experimental Gardens.

A sponge soaked in 10% honey water was provided as an artificial nectar source for females for at least 24 hours before placement into cages. Cages were secured with tent pegs to prevent escape, and a Tinytag datalogger was also placed within each of five cages within this area (at each corner and close to the centre). Females remained in the cages for at least two full days of warm, sunny weather after which point butterflies were collected and plants were removed to be inspected for eggs. Eggs were counted by removing leaves or leaf fragments into labelled petri dishes, ensuring all eggs were collected to avoid double-counting in future assays. Each plant was cross-checked by a second observer to confirm egg counts, and the count repeated if >5% variation in numbers was observed. Host preference was quantified as the number of eggs laid per plant species per female. All plants were given at least 12 hours to recover before being used again for host preference assays. This reuse of plants allowed tests for significant variation across plants in their attractiveness across all females (Results: Oviposition preference varies among individuals and sites and shows high repeatability and is robust to environmental variation). All data were recorded with corresponding site codes, plant IDs, and timestamps. Assays of 41 individuals were repeated (once or twice) to confirm consistency in host preference across time and temperature variation.

### 2023 assays of host preference

To test for changes in host preference over time, and for the effects of host plant preference on larval development under common garden conditions, 69 *A. agestis* females were also collected from second generation individuals from August – September of 2023 (20 generations later). Protocols used were identical to those in 2013, although here the focus was on collecting a range of females that differed in preference profile, in order to test for the effects of parental preference on offspring fitness across host plants. Females were collected from three sites across the UK in 2023, including the two long-established sites (present in 1970–1982): Swyncombe Downs (38 females) and Beacon Hill (8 females), and a newly colonized habitat (since 1995–1997), Barnack Hills and Holes (33 females). Each site was visited either once (Beacon Hill) or twice (Swyncombe Downs and Barnack Hills and Holes). As in 2013, collected females were kept in collapsible cages and fed artificial nectar (a 10% mixture of organic honey to distilled water) for ∼ 24 hours following collection. Individual females were then placed into one of 20, 0.5 × 0.5m wire cages (upturned shopping baskets) covered with mesh. Each cage contained four greenhouse-grown plants: two *Helianthemum nummularium* (Rockrose) and two *Geranium molle* using the sample protocols described above, however all plants were grown at the top of the Darwin Building, Gower St, London, WC1E 6BT, UK. Cages were randomly arranged in two rows (∼ 3m * 12m) at the study site (located at 1 Pool Street, UCL East, London, E20 2AF, UK). Host preference was then recorded using the sample protocols described above for 2013. Assays of 10 individuals were repeated (once or twice) to confirm consistency in host preference across time and temperature variation.

### Rearing of eggs collected during 2023 assays

Once eggs were counted, and maternal host preference had been estimated, we then tested the effects of maternal host preference on offspring performance on either host plant (*Helianthemum nummularium* or *G.molle*). A makeup brush was used to brush eggs into 90mm petri dishes containing dampened filter paper. Eggs from each mother were then pooled together and left to develop in a CT room at 20°C with a light:dark cycle of 16:8, until larvae had hatched from these eggs. Of the 60 females from which host preference data was recorded, 49 had sufficient eggs for larval rearing assays (≥ 8), with the mean number of eggs being 63. After ∼7 days, larvae from each family were collected and randomly assigned to their respective dietary treatment (*Helianthemum nummularium* (Rockrose) or *Geranium molle)*. Fresh leaves were taken from the same greenhouse grown plants described in the 2013 host preference assay methods. All larvae were reared in 90mm petri dishes, with a maximum initial density of 12 larvae per dish to avoid overcrowding.

All larvae were reared under a diurnal variation in temperature to more accurately mimic natural temperature (Brakefield and Mazzotta, 1995; Brakefield and Kesbeke, 1997; Colinet *et al*., 2007; Bozinovic *et al*., 2011; Zanco *et al*., 2025). Larvae (a total of 1441 across 49 females) were reared in a temperature cabinet under fluctuating thermal conditions of 19 ± 3°C, with a light:dark cycle of 16:8 to ensure direct development (Burke *et al*., 2005) and a relative humidity of 70%. Rearing was undertaken in four temporal blocks (starting on 1/9/2023, 3/9/2023, 7/9/2023 and 11/9/2023) determined by when mothers were collected and tested for host preference over the summer. Leaves and filter paper were changed either every second day or daily during heavy feeding periods to minimise infection. Larvae were staged every ∼4 days until 50% of larvae reached instar five. At instar five, if any petri dishes contained more than five larvae it was split across more petri dishes to avoid cannibalism. Each petri dish was checked daily for pupae once larvae began to enter instar stage five. Once pupation occurred, pupae were removed from petri dishes and placed into glass *Drosophila* vials with a diameter of 25mm (1 pupae per vial), plugged with cotton wool, and returned to the same growth chamber to develop (19 ± 3°C and a light:dark cycle of 16:8).

### Development Time, Viability and Metabolic rate

Vials were checked daily for newly emerged adults. Development time to adult or pupae was determined based on how long it took for adults to emerge from the date at which instar one larvae were allocated to experimental treatments. Eggs were considered viable if they pupated. Once adults emerged, a piece of filter paper soaked in 10% honey solution was placed in the vial for no more than 72 hrs post emergence, at which point all adults had their resting metabolic rate recorded using the standard protocol recommended for MAVEn™ System for Small Insect Flow-Through Respirometry. The respirometer lights were turned off and the temperature was set to 25 °C. Dried and CO2 free air was pushed at a rate of 30 ml/sec through the sealed chambers. Each of the 228 butterflies that emerged from pupae were allowed at least 10 minutes in the respirometer before measurements commenced. Air was let to flow through each chamber for 2 minutes before the chamber was sealed again. This cycled through all active chambers and one empty chamber, controlling for drifts in the gas measurement, for ∼ 5 cycles. The amount of CO2 expelled from each butterfly and their activity level was recorded by a LICOR850 machine utilising infrared red radiation to record locomotor activity. Data from the first two cycles to ensure resting metabolic rate was recorded. Time spent in the respirometer, as well as the position of the chamber were used as variables in the subsequent statistical analyses. After the respirometry was concluded, the butterflies were knocked out using CO2 and weighed to the nearest 0.01 mg on a Sartorius ME 5 microbalance (sensitivity ± 1 µg). Mass-independent metabolic rate was calculated by adjusting CO production for adult weight using a linear regression model.

### Wing Size

Wing centroid size was used as a proxy for final body size (Clemson, Sgrò and Telonis-Scott, 2016). All butterflies were dissected after respirometry. Both the left and right wings of 344 butterflies were mounted using double sided tape on glass slides and the torsos were frozen at -20°C (Supplementary Figure 1a). The left forewing was used to measure wing length and size. All wings were photographed using a Leica M80 stereo microscope (Leica, Heer-brugg, Switzerland). At the start and end of photographing, as well as every 30 photos taken, a 10mm scale was photographed to ensure that the scale was known and did not change. The scales were compared in tpsDIG using the protocol outlined in Clemson et al. (2016). The sex was determined and wing centroid size was measured using the software tpsDIG and CoordGen8 (Clemson, Sgrò and Telonis-Scott, 2016). To develop suitable landmarks and test repeatability a standard protocol developed for *Drosophila* landmarking by Clemson et al. (2016) was used in conjunction with existent examples from papers investigating Lepidoptera species (Benítez *et al*., 2011; Bereczki *et al*., 2014; Solonkin *et al*., 2021). A total of eight landmarks were selected (see Supplementary Figure 1b) that consistently produced an R score over 0.99 when repeated measurements were made on 40 wing pairs by two investigators. The coordinates were processed by CoordGen8 to obtain the centroid size in mm.

### Quality analyses of RR and GM host plants at 19 vs 24°C and under well-watered vs drought conditions

A pooled 10 g sample of either all Rockrose or all *Geranium molle* leaves from all plants used for host preference assays in 2023 was submitted to Alex Stewart Agriculture Ltd (21 Sefton Business Park, Olympic Way, Netherton, Liverpool, Merseyside, L30 1RD, United Kingdom) for nutritional analysis. The laboratory is UKAS accredited (ISO/IEC 17025:2017) and routinely applies validated protocols for food and agricultural testing. Protein content was quantified using the Dumas combustion method (07L.1.23), which involves complete combustion of the sample and measurement of nitrogen released to estimate protein concentration, providing a widely accepted alternative to the Kjeldahl method. Available carbohydrate content was measured using method 07L.1.19, which estimates the proportion of digestible carbohydrates by enzymatically breaking down polysaccharides and oligosaccharides into constituent sugars before quantification. This approach provides an assessment of the fraction of carbohydrate available as an energy source, excluding indigestible fibre.

To assess the effects of both drought and heat stress on the growth and nutritional content of each plant species, 20 *Helianthemum nummularium (*Rockrose) plugs (purchased from Naturescape (address above)) or 20 *Geranium molle* seeds (purchased from Emorsgate Seeds (address above)) were grown in Sinclair Professional Multi-Purpose Compost (Firth Road, Lincoln, LN6 7AH, United Kingdom) under a fully crossed, pairwise design: either 19 ±3°C or 24 ±3°C, and under well-watered (every 2–3 days) or drought (once per week) conditions. All plants were kept in growth chambers which were set to a light:dark cycle of 16:8. Physiological traits were measured using a Dualex optical sensor (DX18068), Pessl Instruments, which provides non-destructive measurements of chlorophyll, flavanoids and anthocyanins. For each plant, three measurements were taken from both sides of two fully expanded leaves, to estimate within-plant vs among plant as well as among species variation.

### Statistical analysis

All statistical analyses were conducted in R (version 2024.12.0+467), using the lme4, lmerTest, emmeans, and boot packages. Type III ANOVA tables were generated using the lmerTest package to evaluate the significance of model terms. Pairwise comparisons of estimated marginal means were conducted using the emmeans package, with p-values adjusted for multiple comparisons. Summary statistics for pairwise comparisons (means and standard errors) were calculated using the dplyr package to visualise treatment effects.

### Host preference

The reliability of host preference estimates for each mother increases with number of eggs laid, and previous studies (e.g. Buckley and Bridle 2014) show consistently higher egg numbers on *Geranium molle* plants. To test for within site variation and temporal variation in host plant use, we employed permutation tests (10,000 iterations) and bootstrapped confidence intervals for egg allocation and preference scores. One-sample t-tests were used to compare observed egg allocation to null expectations of random oviposition to determine if mothers were more likely to lay on the dominant host plant growing where they were collected from in 2013. To assess the consistency of maternal host preference within individual mothers, we calculated repeatability using likelihood ratio tests comparing models with and without individual identity as a random effect. 2013 and 2023 data was combined, so that year was included as a fixed effect to account for temporal variation. Generalized linear mixed models (GLMMs) were used to evaluate all other effects on maternal host preference, with fixed effects including year, site, and host plant identity, and random effects including individual plant identity and mother ID.

### Larval traits

Linear mixed-effects models (LMMs) were used (lme4 package) to assess the effects of diet, maternal preference, and adult sex on larval performance traits (viability, development time, wing size, and mass-independent metabolic rate), with mother and petri dish included as random effects, except for viability, which was measured per petri dish so only mother was included as a random effect. Mass-independent metabolic rate was calculated by adjusting CO production for adult weight using a linear regression model.

### Plant traits

LMMs were also used to test the effects of temperature and water supply on *Geranium molle* and Rockrose (traits measured included: chlorophyll, flavonoids and anthocyanins), with plant ID as a random effect.

## Results

### Oviposition preference varies among individuals and sites, and shows high repeatability

To test to what extent our assays of host preference variance among and within mothers reflect among genotype variation, we first tested whether maternal host preference or individual-level variation was influenced by ambient temperature in both the 2013 and 2023 datasets. Oviposition preference for Rockrose relative to *Geranium molle* was not significantly associated with ambient temperature in cages during assays (2013: p = 0.555; 2023: p = 0.458), suggesting that temperature variation during trials did not influence maternal host choice. Next, we quantified the repeatability of host preference scores across 41 and 10 mothers within 2013 and 2023 respectively to assess the consistency of individual females’ host choices. We found that maternal preference was highly repeatable (repeatability = 0.76, 95% CI (0.61, 0.85); likelihood ratio test: D = 42, p < 0.001), demonstrating that – even with relatively low numbers of repeats - most variation in preference was explained by consistent differences in host preference among females rather than by within-female variability over time.

### Oviposition preference and is robust to environmental variation

To assess whether variation among individual plants within host plant species influenced host preference estimates, we compared GLMMs with and without individual plant identity as a random effect. While including plant identity increased uncertainty around preference estimates, the direction and magnitude of maternal preference remained consistent across models (2013: –0.22 ± 0.02 SE without plant ID; –0.38 ± 0.31 SE with plant ID; 2023: –0.77 ± 0.04 SE without plant ID; –0.85 ± 0.24 SE with plant ID). These results indicate that although individual plants within species differ in attractiveness to these mothers, even when grown in a common greenhouse and from commercial seed stock, this variation does not systematically bias detection of host preference. Therefore, plant identity was excluded from subsequent host preference models, allowing us to model each mother’s preference averaged across all four plants she was offered.

The evidence from these analyses supports the conclusion that our short-term assays of maternal host preference in *A. agestis* provide an estimate of a consistent behavioural trait that is significantly repeatable despite short-term environmental variation or individual plant differences. The strong repeatability and consistent preference signals across multiple years reinforce the biological relevance of host choice in shaping host preference patterns. This foundation enables us to confidently proceed with analyses of longer-term behavioural shifts and evolutionary dynamics in host use.

### Host plant preference shows differences in mean and reduced variation in the recently-expanded range

To test how host preference behaviour varies across the UK range of *A. agestis*, we first quantified host use by bootstrapping egg counts across host types (10,000 randomisations). This approach provided robust estimates of per-mother and site-level egg allocation, along with confidence intervals, accounting for variation in sample sizes and individual egg numbers (Figure 1a,b). A permutation test then confirmed significant variation in mean Rockrose use across sites (F = 4.72, p < 0.001), but no significant difference in mean Rockrose use between old and new sites (p = 0.65), suggesting that colonisation history does not explain spatial variation in host use (Figure 1d).

To assess whether females preferentially oviposited on the locally dominant host, we compared the mean egg allocation of individual females to a null expectation using a one-sample t-test. Mothers laid significantly more eggs on the dominant host than expected by chance (t = 3.60, p < 0.001, Figure 1d), indicating that local plant availability strongly influences the frequency of host preference observed at a given site. We then tested whether colonisation history also affects this pattern using a linear mixed-effects model. Mothers from newly colonised sites laid a 16% greater proportion of their eggs on the dominant host compared to those from established sites (estimate = –0.16, p = 0.037, Figure1d).

**Figure 1.**
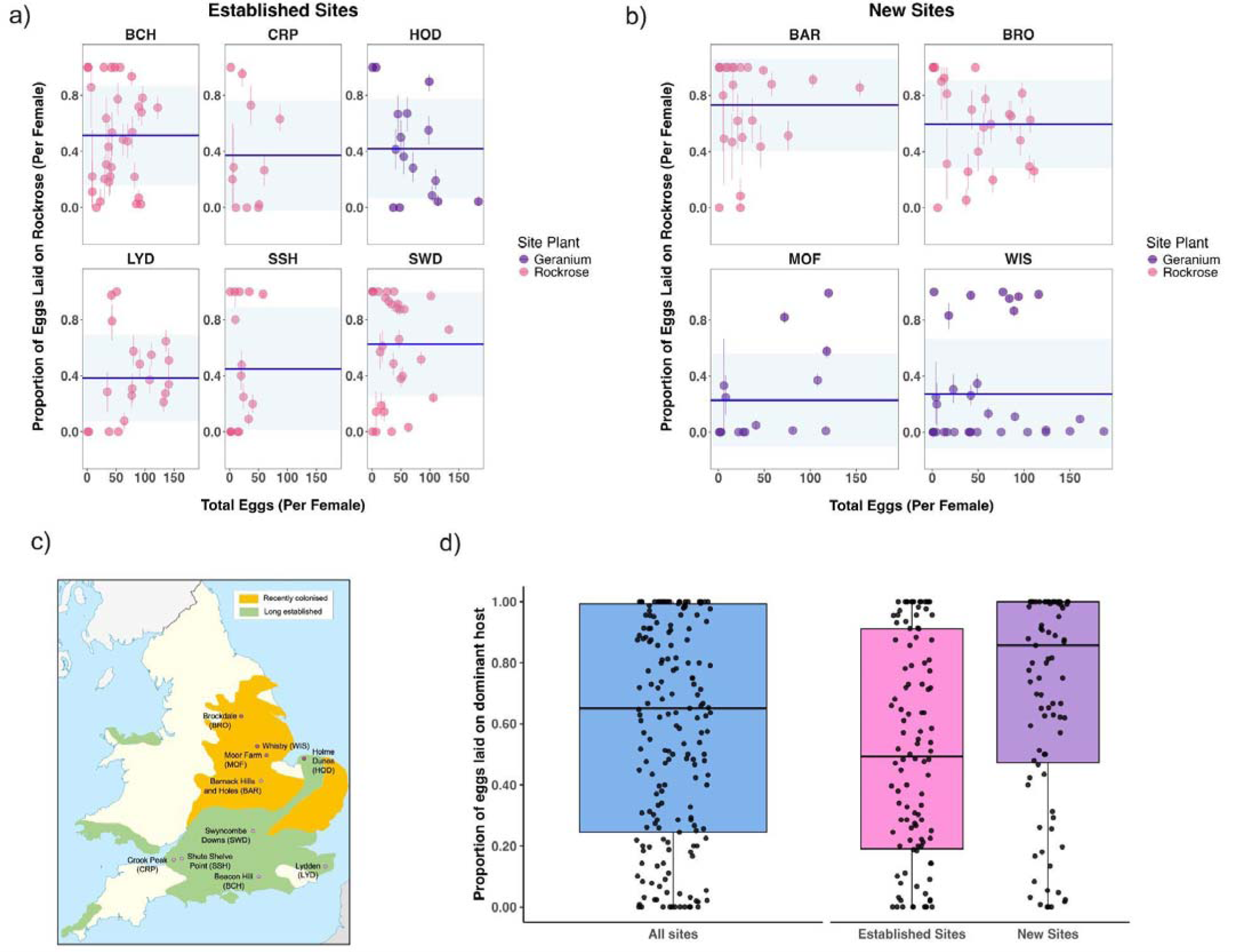
In newly colonised sites, mothers are more likely to specialise on the locally dominant host plant, while in established sites, mothers show broader host-plant preferences regardless of local plant availability in 2013. The average number of eggs laid per host plant was calculated for each mother, with 95% confidence intervals estimated using 10,000 bootstrapped resamples (a,b). Each mother’s preference was used to calculate the mean and standard deviation in preference at each site, shown as a blue line and ribbon (a,b). Map of sites sampled in 2013, classified by colonisation history (established or newly colonised) and dominant host plant family (Geranium molle (purple) or Rockrose (pink)) (c). We tested whether maternal preference varied across sites using 10,000 permutations and found significant spatial variation in host preference (F = 4.72, p < 0.001) (a,b,c). The boxplots display the distribution of maternal preferences, with the horizontal line indicating the median (d). Mothers laid a greater proportion of eggs on the locally dominant host than expected by chance (t = 3.60, p < 0.001) (d). A linear mixed effects model showed that mothers from established sites laid a significantly lower proportion of their eggs on the dominant host compared to those from newly colonised sites (estimate = -0.16, p = 0.037), indicating stronger host specialisation in the new range (d).

### Maternal preference for Geranium molle has increased over a decade, especially in historically Rockrose-dominated populations

In 2023, we returned to three sites: one newly colonised - Beacon Hill (BCH), and two established - Barnack Hills and Holes (BAR) and Swyncombe Downs (SWD) (Figure 2c), where 20 *A.agestis* generations earlier, maternal preference was either skewed towards Rockrose or split evenly between the two host plants (Figure 2a,b). First, we compared patterns of host use between 2013 and 2023 (Figure 2a,b). A linear mixed-effects model including year as a fixed effect and site as a random effect revealed a significant temporal shift in host preference (Figure 2d). In 2023, females laid 24% fewer eggs on Rockrose compared to 2013 (estimate = –0.241, *t* = –4.38, *p* < 0.001) (Figure 2d). To assess whether the trajectory of this shift varied spatially, a second model was constructed with site, year, and their interaction as fixed effects. The decline in Rockrose preference remained significant (estimate = –0.328, *t* = –3.62, *p* < 0.001), with no significant main effects of site, or their interaction terms. These results demonstrate consistent reduction in maternal preference for Rockrose across the sampled landscape.

**Figure 2.**
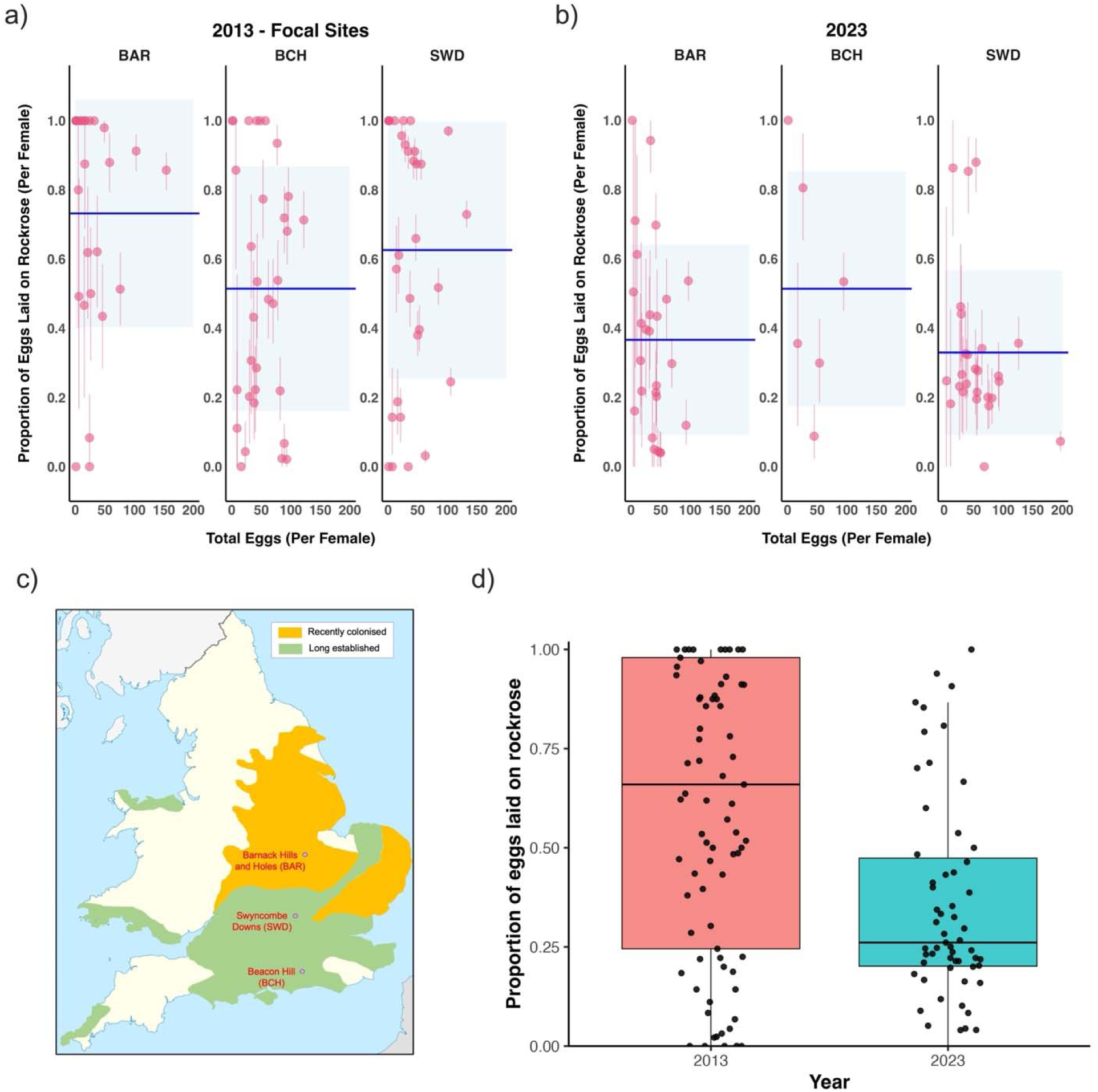
Maternal preference for host plants has shifted over time, with significantly more eggs laid on Geranium molle in 2023 than in 2013. The average number of eggs laid per host plant for each mother was calculated, with 95% confidence intervals estimated using 10,000 bootstrapped resamples (a,b). Each mother’s preference was used to calculate the mean and standard deviation in preference at each site (shown as a blue line and ribbon) (a,b). Map of the three sites sampled in both 2013 and 2023: one newly colonised (BCH) and two established (BAR and SWD), all of which were historically associated with Rockrose (c). The boxplots display the distribution of maternal preferences, with the horizontal line indicating the median (d). A linear mixed-effects model revealed a significant decline in the proportion of eggs laid on Rockrose between 2013 and 2023 (estimate = –0.241, t = –4.38, p < 0.001) (d). Host preference data was gathered from 60 females whose eggs were used for larval rearing experiments and phenotyping (see Figure 3 and 4).

### Geranium molle use in laboratory conditions enhances larval performance regardless of maternal host preference

We next tested the preference–performance hypothesis by assessing how larval and adult performance traits are influenced by host plant diet and maternal host preference. 1412 larvae from 49 maternal families with known host preference (Figure 2b,d) were evenly split and reared on either Rockrose or *Geranium molle*, effectively reciprocally transplanting eggs evenly among host plants regardless of their maternal preference. We assessed four key fitness traits: survival to pupation, development time, adult body size (wing centroid size), and resting mass-independent metabolic rate (MIMR, measured while resting). Across all traits, maternal host preference had no significant effect when including maternal identity and, where applicable, petri dish as random effects: development time (estimate = -0.91 ± 1.18, χ² = 0.60, p = 0.44), survival to pupation (estimate = 0.047 ± 0.077, χ² = 0.38, p = 0.54), wing size (estimate = 0.11 ± 0.25, χ² = 0.21, p = 0.65), and resting MIMR (estimate = -0.00003 ± 0.00005, χ² = 0.40, p = 0.53). This finding suggests that even if a mother oviposited predominantly on Rockrose, or evenly across both hosts, offspring still performed the same on *Geranium molle* as offspring from mothers that oviposited predominantly on *Geranium molle.* Thus, maternal preference did not influence offspring performance under laboratory conditions with ample food (Figure 3).

**Figure 3.**
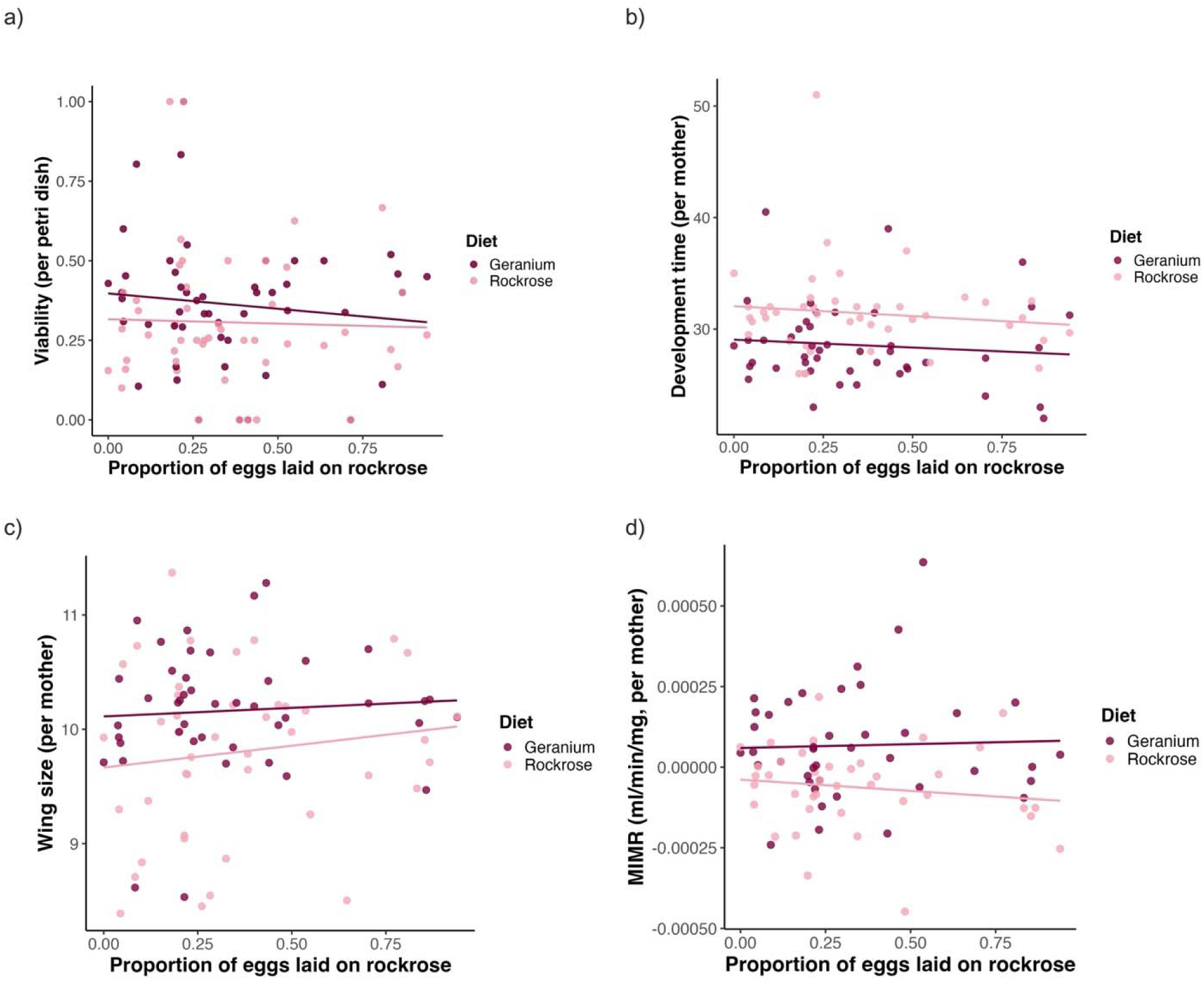
Family-level performance of Brown Argus offspring in response to larval diet. Each panel shows the mean value of offspring from each mother for a key fitness trait: (a) viability to pupation, (b) larval development time, (c) adult wing size (centroid size), and (d) resting mass-independent metabolic rate (MIMR). Maternal families are indicated by points, with offspring split between rearing on Geranium molle (purple, “Geranium”) or Rockrose (pink, “Rockrose”). Regression lines for each diet are fitted separately to illustrate the continuous relationship between maternal host preference (proportion of eggs laid on Rockrose) and offspring performance. Mixed-effects models including mother identity (and Petri dish for all traits other than viability) as random effects revealed that maternal host preference had no significant effect on any trait (development time: χ² = 0.60, p = 0.44; viability: χ² = 0.38, p = 0.54; wing size: χ² = 0.21, p = 0.65; MIMR: χ² = 0.40, p = 0.53).

### *Geranium molle* enhances performance without detectable trade-offs

Larvae reared on *Geranium molle* consistently outperformed those reared on Rockrose across multiple traits (Figure 4a-d). Survival from egg to pupae was 6.9% higher on Geranium compared to Rockrose (p = 0.027), with significant variation among mothers (Mother random effect: variance = 0.007, p = 0.03), indicating genetic or maternal effects. Development time was 2.61 days longer on Rockrose than on Geranium (p < 0.001), also influenced by mother identity (variance = 1.324, p < 0.05). Adult wing size was 0.43 mm smaller on Rockrose relative to Geranium (p < 0.001), with significant mother-level variation (variance = 0.146, p < 0.05). Resting mass-independent metabolic rate (MIMR) was 1.6 × 10 ml/min/mg higher on Geranium (p = 0.0019) and showed no detectable variation among mothers (variance ≈ 0), suggesting this trait is largely environmentally driven under laboratory conditions. Development time did not explain variation in adult size when modelled alongside diet (p = 0.95). To assess potential trade-offs, correlations among development time, wing size, and resting MIMR were examined. Correlations were weak and did not differ significantly between diets (all p > 0.05), indicating that larvae exploit *Geranium molle* without apparent physiological compromises. Sex-specific diet interactions were also non-significant (all p > 0.05), showing that these diet effects are consistent across males and females.

**Figure 4.**
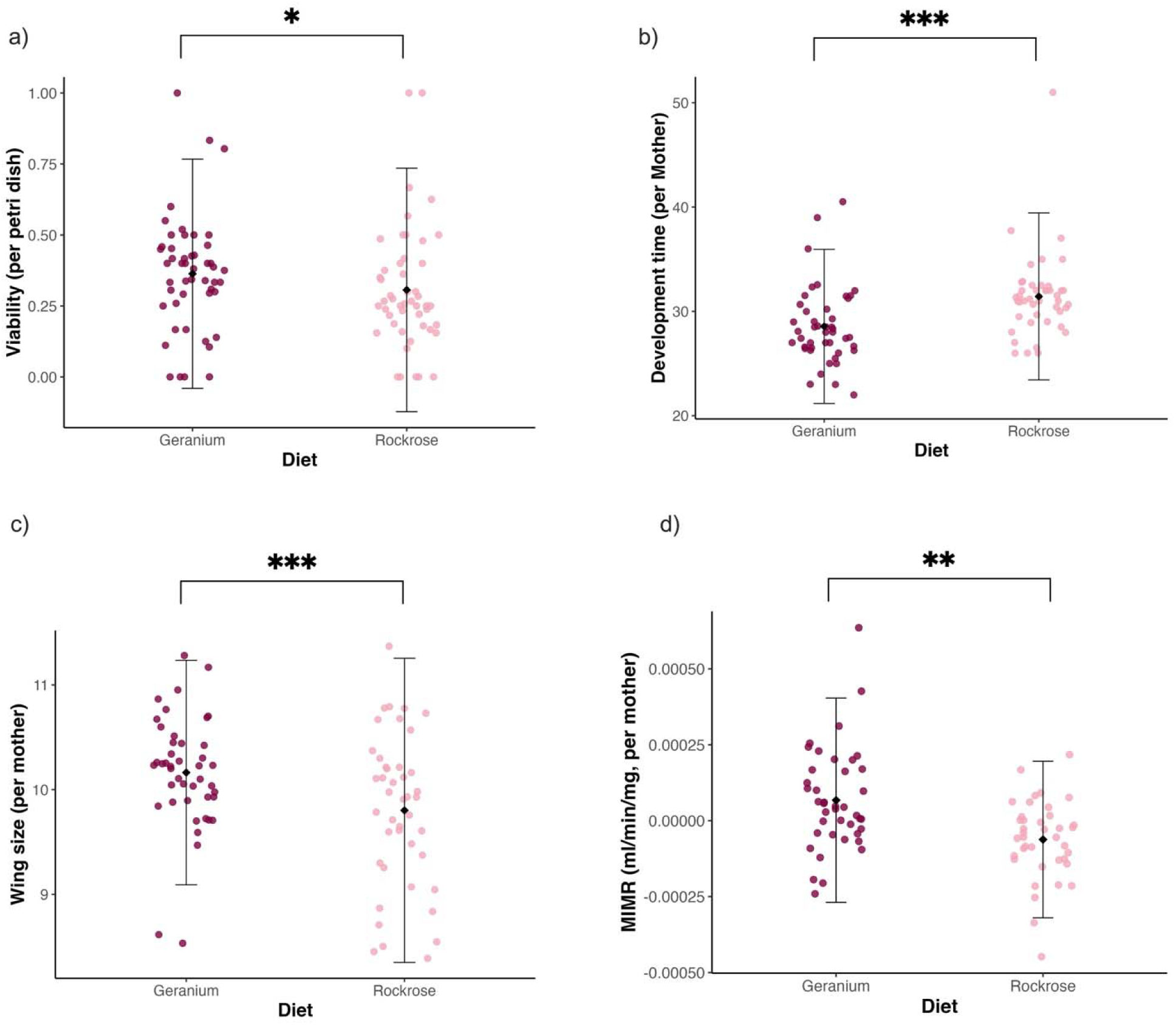
Larvae reared on Geranium molle showed higher viability, faster development, larger adult body size, and elevated resting mass-independent metabolic rates (MIMR) compared to those reared on Rockrose. Survival to pupation was significantly higher on Geranium molle (p = 0.027; mean difference = 6.9%) and showed significant variation among mothers (Mother random effect: variance = 0.007, p = 0.03) (a). Larvae developed faster on Geranium molle, with development time on Rockrose on average 2.61 days longer (p < 0.001), and development time was also influenced by mother identity (variance = 1.324, p < 0.05) (b). Wing centroid size, used as a proxy for adult body size, was greater for adults reared on Geranium molle (p < 0.001) and showed significant mother-level variation (variance = 0.146, p < 0.05) (c). MIMR, calculated by adjusting COC production for adult weight using a linear regression model, was significantly higher in adults reared on Geranium molle (mean difference ≈ 1.6 × 10CC ml/min/mg, p = 0.0019), but showed no detectable variation among mothers (variance ≈ 0), suggesting this trait is largely environmentally driven (d). Each point represents the mean per mother (or per Petri dish for viability), with black lines indicating the mean and standard deviation. All plots display mean ± SD. Development time did not explain variation in adult size when modelled alongside diet (p = 0.95). Statistical significance is indicated by asterisks: : p < 0.05, *p < 0.01, and **p < 0.001***.

### Rockrose is more resilient to heat and drought stress than Geranium molle

To investigate drivers of host selection and performance, we analysed the macronutrient composition and caloric content of the leaves of plants that were fed to larvae. We found that *Geranium molle* had a higher P:C ratio (0.53) than Rockrose (0.44), despite having lower overall caloric content (Table 1). This pattern suggests that *Geranium molle* provides a relatively protein-rich diet, but that larvae must consume more of it to obtain the same energy value as Rockrose.

We next examined how two temperature regimes (19 ± 3°C and 24 ± 3°C) and two water treatments (well-watered and drought) influenced secondary metabolites in Rockrose and *Geranium molle*. Chlorophyll, a proxy for photosynthetic potential, was strongly species-specific and temperature-sensitive (Figure 5a). Across all treatments, Rockrose consistently contained more chlorophyll than *Geranium molle* (dry 19°C: RR 27.42 vs GM 16.47; wet 19°C: RR 30.36 vs GM 15.54; all P < 0.001; Figure 5a). Elevated temperature reduced chlorophyll in *Geranium molle* under dry conditions (16.47 at 19°C vs 13.56 at 24°C; P = 0.005), whereas Rockrose remained largely unaffected (Figure 5a). Drought alone had no significant effect (all P > 0.2). Anthocyanins, which act as pigments and stress-inducible antioxidants, varied with species, temperature, and water availability (Figure 5b). *Geranium molle* had higher anthocyanin levels than Rockrose under drought conditions (dry 19°C: GM 0.160 vs RR 0.104; P < 0.001; Figure 5b). In Rockrose, warming under wet conditions increased anthocyanins (0.118 at 19°C vs 0.153 at 24°C; P < 0.001), whereas *Geranium molle* decreased under dry warming (0.160 at 19°C vs 0.146 at 24°C; P = 0.014; Figure 5b). Drought responses were species and temperature-dependent: at 24°C, drought reduced anthocyanins in Rockrose (0.153 vs 0.128; P = 0.028), while other contrasts were non-significant (Figure 5b). Flavonoids, which function primarily as UV screens and constitutive antioxidants, showed consistently higher levels in *Geranium molle* than Rockrose (dry 19°C: GM 0.276 vs RR 0.084; wet 19°C: GM 0.167 vs RR 0.058; all P < 0.01; Figure 5c). In Rockrose, drought induced a significant increase at 19°C (0.167 wet vs 0.239 dry; P = 0.018), but temperature had no effect (Figure 5c). By contrast, *Geranium molle* maintained constitutively high levels regardless of treatment (Figure 5c).

**Table 1.** Nutritional composition and caloric content of Rockrose and Geranium molle leaves. Rockrose had a lower P:C ratio (0.44) and higher caloric content (73.36 kcal) compared to Geranium molle (P:C ratio of 0.53 and caloric content of 47.28 kcal).

| Plant species | Protein (%) | Carbohydrate (%) | P:C Ratio | Caloric Content (kcal) |
| --- | --- | --- | --- | --- |
| Rockrose | 5.52% | 12.52% | 0.44 | 73.36 |
| *Geranium molle* | 4.09% | 7.72% | 0.53 | 47.28 |

**Figure 5.**
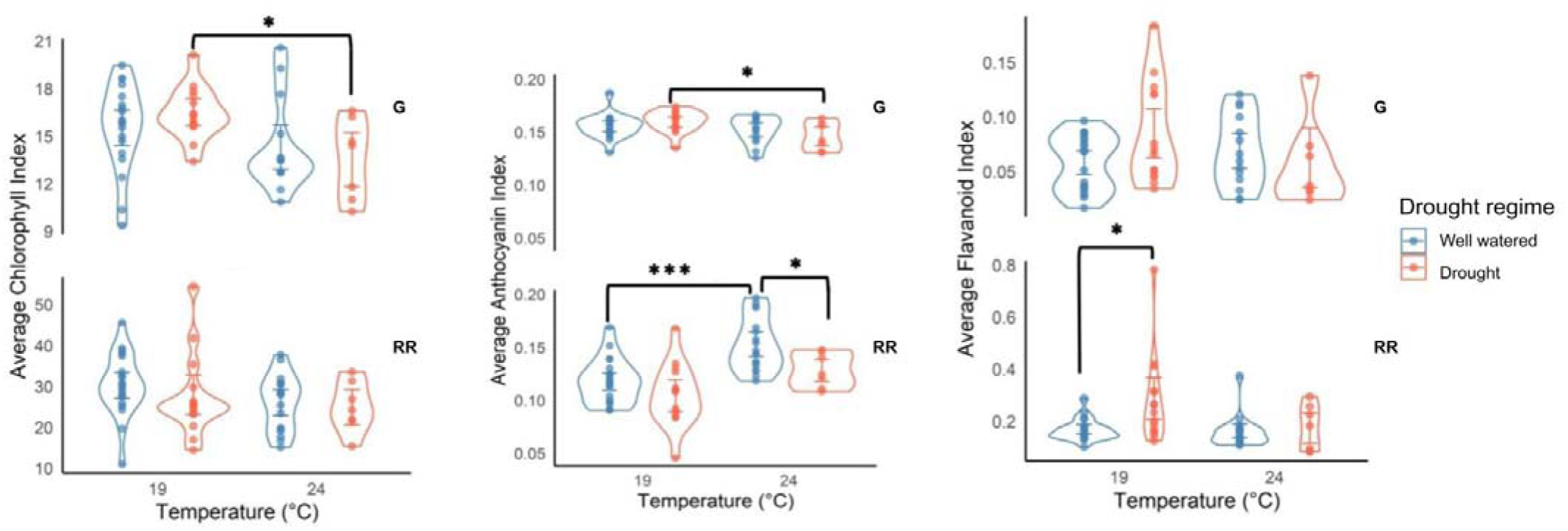
Panels show (a) chlorophyll index, (b) anthocyanin index, and (c) flavonoid index across two host species (Rockrose and Geranium molle) under factorial combinations of temperature (19 ±C3°C and 24 ±C3°C) and water availability (Well-watered and Drought). Points represent individual leaves, with violin distributions overlaid. Chlorophyll content (a), was strongly species-specific and temperature-sensitive. Across all conditions, Rockrose had higher chlorophyll than Geranium molle (dry 19°C: 27.42 vs 16.47; wet 19°C: 30.36 vs 15.54; all P < 0.001). Elevated temperature reduced chlorophyll in Geranium molle under dry conditions, from 16.47 at 19°C to 13.56 at 24°C (P = 0.005), whereas Rockrose levels were largely unchanged. Drought alone had no significant effect (all P > 0.05). Anthocyanin levels (b) varied with species, temperature, and drought. Geranium molle had more anthocyanis than Rockrose under dry conditions (dry 19°C: 0.160 vs 0.104, P < 0.001). Temperature increased anthocyanins in Rockrose under wet conditions, from 0.118 at 19°C to 0.153 at 24°C (P < 0.001), while Geranium molle decreased under dry warming, from 0.160 at 19°C to 0.146 at 24°C (P = 0.014). Drought effects were species- and temperature-dependent: at 24°C, drought reduced anthocyanin levels in Rockrose from 0.153 to 0.128 (P = 0.028), whereas other combinations were non-significant. Flavonoid levels (c) were strongly species-specific, with higher constitutive levels in Geranium molle (dry 19°C: 0.276 vs 0.084; wet 19°C: 0.167 vs 0.058; all P < 0.01). Drought significantly increased flavonoids in Rockrose at 19°C, from 0.167 under wet conditions to 0.239 under dry conditions (P = 0.018), whereas temperature effects were non-significant. Statistical significance is indicated by asterisks: p < 0.05, *p < 0.01, and **p < 0.001***.

## Discussion

In this study, we estimated maternal host preference, larval performance, and host plant trait data to understand the ecological and evolutionary dynamics of host use in *A. agestis* and infer the consequences of the evolution of new biotic interactions for population persistence under climate change. We show for the first time that: (i) maternal host preference varies both within and among sites in *A.agestis*, and that *Geranium molle* preference has increased significantly between 2013 and 2023, even in historically Rockrose-dominated populations; (ii) maternal host preference is highly repeatable among females but does not predict offspring performance; (iii) larvae reared on *Geranium molle* consistently outperform those on Rockrose across multiple traits, without detectable trade-offs; (v) *Geranium molle* has a higher P:C ratio than Rockrose but is less calorie dense, suggesting that its apparent advantages under well-fed laboratory conditions may not translate directly to resource-limited environments; and (vi) *Geranium molle* is more vulnerable than Rockrose to heat and drought stress, raising the possibility of ecological traps.

### Host preference is shifting across the range, not just at the expansion front

We show that across the UK, host preference in *A. agestis* is strongly spatially structured: females preferentially lay eggs on the locally dominant host. This pattern occurs in both newly colonised and long-established populations, but its influence is stronger in the north, where *Geranium molle* has become increasingly abundant while Rockrose is declining (Pateman *et al*., 2012; Bridle *et al*., 2013; Buckley and Bridle, 2014). In these northern populations, females may specialise on *Geranium molle* simply because it is both more available, and increasing in availability (Pateman *et al*., 2012; Bridle *et al*., 2013; Buckley and Bridle, 2014), raising the possibility that Rockrose-associated populations could contract or disappear over time. These data align with similar patterns observed in *M. cinxia*, whereby local host abundance drives preference (Kuussaari, Singer and Hanski, 2000).

In addition to these spatial differences, we observed a strong temporal trend: between 2013 and 2023, Rockrose use declined consistently across sites, including long-established populations where it had previously dominated. This indicates that host preference is shifting towards *Geranium molle* range-wide, pointing to evolutionary change that is not confined to colonising fronts. This pattern is consistent with evidence that gene flow is widespread throughout *A. agestis*’ range (de Jong *et al*., 2023), and highlights how core populations likely play an important role in driving and sustaining long-term stability in new populations, while also preventing contractions (Thomas *et al*., 2001; Lenoir and Svenning, 2015; Saastamoinen *et al*., 2018; Hällfors *et al*., 2024).

### Range-wide specialisation challenges assumptions about diet breadth and expansion

Broad dietary breadth is generally thought to facilitate range expansion, allowing populations to exploit novel or abundant hosts (Lancaster, 2020; Sunde et al., 2023). This pattern is observed in butterflies such as the Edith’s Checkerspot and *P. mannii*, where populations with broader diets more successfully colonise new habitats (Neu *et al*., 2021; Singer and Parmesan, 2021). By contrast, *A. agestis* is narrowing its host preference, increasingly specialising on *Geranium molle*. Crucially, this specialisation is occurring not only at the expansion front, but also in long-established populations; thus demonstrating that rapid evolutionary changes in host use are spreading throughout the species’ range. This divergence from general predictions has important implications. First, it shows that assumptions about diet breadth during range shifts do not always hold in specialist systems. Second, it highlights potential evolutionary risks: tight coupling to a single host may increase vulnerability to host instability under climate change.

This range-wide specialisation may also have long-term evolutionary consequences. Rapid adaptation can facilitate persistence and support colonisation of new environments, yet strong directional selection on host-use traits can erode standing genetic variation, narrowing the genetic base available for future responses as environments become more unpredictable (Hoffmann and Bridle, 2022). *A. agestis* therefore demonstrates a paradox some species might be expected to experience under ongoing climate change: the same evolutionary shifts that enabled rapid northward expansion and increased reliance on *Geranium molle* may now constrain the species’ long-term adaptive potential.

### Nutritional benefits of Geranium molle may drive rapid evolutionary shifts

We found that larvae consistently perform better when fed *Geranium molle*, under well fed laboratory conditions when compared to Rockrose. These data are in line with findings previously reported by Bodsworth (2002) who found that larvae reared on *Geranium molle* developed faster and were larger than those reared on Rockrose. It is possible that the P:C ratio of *Geranium molle* compared to Rockrose leaves is a key driver of these effects. Indeed, substantial research has demonstrated that dietary P:C ratio strongly influences lifetime performance across taxa, with high P:C diets resulting in faster development, increased viability, larger final body sizes and higher reproductive output (e.g. Lee *et al*., 2008; Simpson and Raubenheimer, 2009; Clissold and Simpson, 2015; Lee, 2015; Solon-Biet *et al*., 2015; Helm *et al*., 2017). However, although *Geranium molle* has a higher P:C ratio, its lower absolute levels of protein and carbohydrate suggest that in food-limited years, larvae might not obtain enough calories to sustain rapid development, potentially reducing performance on this host. More broadly, we also know that dietary macronutrient balance interacts with other environmental stressors such as temperature to drive fitness outcomes (Kutz, Sgrò and Mirth, 2019; Chakraborty, Sgrò and Mirth, 2020; Zanco *et al*., 2025), and that micronutrients likely play a significant role in mediating these effects (Zanco *et al*., 2020; Piper *et al*., 2022; Sarmah *et al*., 2025), factors we did not consider in this study.

### Maternal preference does not predict larval success

Despite clear shifts in host preference, maternal preference had no detectable effect on offspring performance under laboratory conditions. Larvae reared on *Geranium molle* developed faster and reached larger adult sizes than those fed on Rockrose, regardless of maternal choice. This indicates that a narrowing of host preference towards Geranium species is unlikely to be driven by precise maternal selection (Buckley and Bridle, 2014). One explanation for the decoupling of female preference and larval performance is that these traits might be genetically and selectively independent (Wiklund, 1975; Thompson, 1988). This could explain why most mothers still laid at least some eggs on Rockrose, even though offspring performed best on *Geranium molle*. Importantly, the persistence of Rockrose use may reflect ecological benefits not captured under well-fed laboratory conditions. In the wild, Rockrose could provide an advantage through its higher caloric availability, particularly in resource-limited years when larvae may be unable to fully capitalise on the higher P:C ratio of *Geranium molle*, as discussed above.

### Diet-driven shifts in adult metabolic rate highlight consequences of host use shifts on adult performance

Resting metabolic rate is the largest component of total energy expenditure in most animals (Speakman and Selman, 2003). Here, we show that larvae fed on *Geranium molle* develop higher mass-independent resting metabolic rates as adults compared to those fed on Rockrose. These are among the first data, alongside Rossi and Niven (2021), to demonstrate lasting effects of larval diet on adult metabolism outside of epidemiological contexts, and in a wild butterfly species. Unlike other traits, metabolic rate was influenced solely by diet, not family background, indicating a direct physiological response to the host plant rather than underlying genetic variation. Elevated adult metabolic rates may carry both costs and benefits: adults reared on *G. molle* not only exhibited higher metabolism but also developed larger wings, which could enhance dispersal ability (Auer *et al*., 2015; Nicholls, Rossi and Niven, 2021; Pocius *et al*., 2022). Indeed, movement rates have been shown to increase in *A.agestis* females that are searching for *Geranium molle*, with individuals spreading more widely across the landscape (Bodsworth et al., 2002). In saying this, increased baseline energy expenditure can reduce survival under resource-limited conditions. Overall, these results underscore how nutritionally induced changes in physiology can have lasting consequences for performance and host exploitation, independent of genetic variation, and suggest that larval diet can shape adult metabolism in ecologically meaningful ways.

### Climate stress increases the risk of ecological traps in increasingly specialised systems

Ecological strategies in plants shape the resources available to herbivores, and for butterflies the nutritional quality and temporal availability of food during the larval stage is strongly influenced by the strategies adopted by their host plants (Dennis *et al*., 2004; Carvajal Acosta, Agrawal and Mooney, 2023). *Geranium molle* and Rockrose exemplify contrasting ecological strategies: *Geranium molle* functions as a ruderal (R; high disturbance, low stress), while Rockrose is a stress-tolerator (S; high stress, low disturbance) (Dennis *et al*., 2004). These contrasting strategies create different consequences for herbivores. Stress-tolerant plants such as Rockrose persist throughout the year but generally provide food of lower nutritional quality. Larval development on these hosts is slower, and dispersal propensity in butterfly populations is reduced (Dennis *et al*., 2004). In contrast, ruderal plants such as *Geranium molle* provide higher-quality food, but only for a limited time. Larvae feeding on these hosts develop more rapidly and populations tend to show greater dispersal capacity (Dennis *et al*., 2004).

A complementary framework comes from the mesic–arid adaptation of host plants. Acosta et al. (2022) demonstrated that herbivores on mesic-adapted plants are vulnerable to drought because these hosts lose vigour under stress, whereas herbivores feeding on arid-adapted plants often maintain, or even improve, performance. In this context, our results show that *Geranium molle*, a mesic-adapted ruderal, suffers pronounced declines in chlorophyll content and photosynthetic activity under heat and drought. Rockrose, by contrast, is both stress-tolerant and arid-adapted: it maintains physiological function under stress but shows stronger inducible increases in secondary metabolites. These features could both explain why most mothers still lay some eggs on Rockrose and be used to predict a future where a narrowing preference for Geranium species could pose a threat to the persistence of *A. agestis*, despite the fitness advantages we recorded under laboratory conditions.

### Conclusions

Our findings show that *A. agestis* is undergoing rapid adaptation in both newly colonised and historically established populations. These findings support a broader view of range shifts as multi-scale processes, where phenotypic and ecological change occurs across the entire range (Thomas *et al*., 2001; Lenoir and Svenning, 2015; Saastamoinen *et al*., 2018; Hällfors *et al*., 2024). By integrating host preference, offspring performance, and plant trait and stress response data, we show how short-term fitness gains on *Geranium molle* can facilitate both persistence and expansion, but potentially at the risk of reduced resilience under climate extremes. Importantly, our results point to both diet quality and availability as a powerful driver of rapid adaptation.

By linking nutritional ecology, physiology, and range dynamics, this study provides a basis for understanding how the rapid evolution of host preference unfolds across species’ ranges and over time. Future work should now link these ecological and physiological patterns to underlying genetic architecture. Approaches such as genome-wide association studies (GWAS) or landscape genomics could identify alleles associated with host preference, performance, or stress tolerance, clarifying how selection operates across the range and how rapidly adaptive traits can spread over space and time (Bridle and Hoffmann, 2022; de Jong *et al*., 2023). More broadly, our results highlight that biodiversity responses to climate change are likely to involve trade-offs: short-term gains in performance and colonisation potential set against long-term vulnerabilities to instability and loss of adaptive capacity.

