## Supplementary Figure 1. for "Does mother know best? Range-wide narrowing of host preference in *Aricia agestis* confers fitness benefits but may incur long-term costs"

**Supplementary Figures**

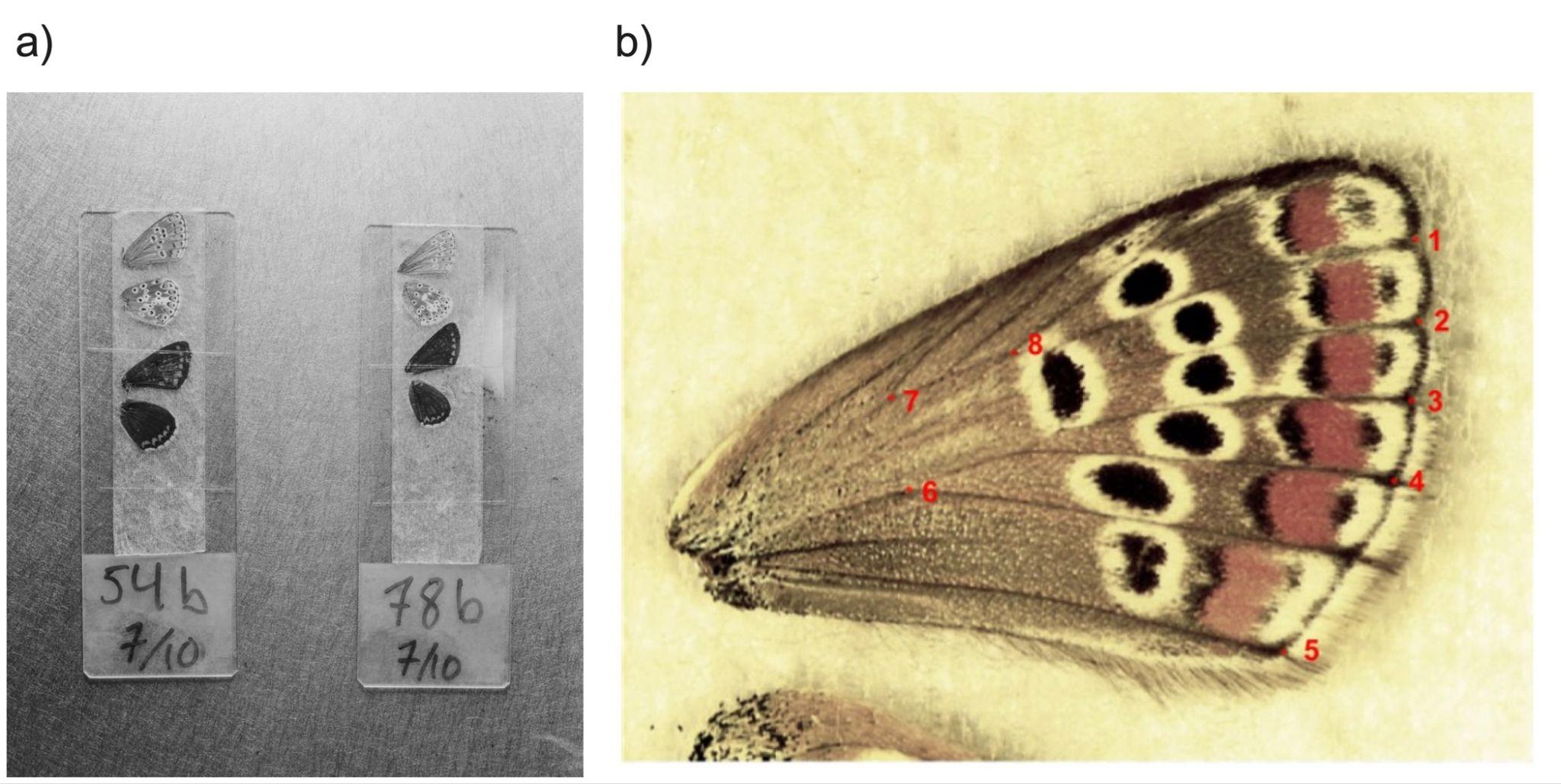

***Supplementary Figure 1.*** *Wings were dissected and placed on glass slides (a). Both the left and right wings of 344 butterflies were mounted using double sided tape (a). A total of 8 landmarks were selected (b) for final wing size analyses that consistently produced an R score over 0.99 when repeated measurements were made on 40 wing pairs (Clemson et al. 2016).*
